# Ten new high-quality genome assemblies for diverse bioenergy sorghum genotypes

**DOI:** 10.1101/2022.09.26.509520

**Authors:** William G. Voelker, Krittika Krishnan, Kapeel Chougule, Louie C. Alexander, Zhenyuan Lu, Andrew Olson, Doreen Ware, Kittikun Songsomboon, Cristian Ponce, Zachary W. Brenton, J. Lucas Boatwright, Elizabeth A. Cooper

## Abstract

Sorghum (*Sorghum bicolor* (L.) Moench) is an agriculturally and economically important staple crop that has immense potential as a bioenergy feedstock due to its relatively high productivity on marginal lands. To capitalize on and further improve sorghum as a potential source of sustainable biofuel, it is essential to understand the genomic mechanisms underlying complex traits related to yield, composition, and environmental adaptations. Expanding on a recently developed mapping population, we generated *de novo* genome assemblies for 10 parental genotypes from this population and identified a comprehensive set of over 24 thousand large structural variants (SVs) and over 10.5 million single nucleotide polymorphisms (SNPs).These resources can be integrated into both ongoing and future mapping and trait discovery for sorghum and its myriad uses including food, feed, bioenergy, and increasingly as a carbon dioxide removal mechanism. We show that SVs and nonsynonymous SNPs are enriched in different gene categories, emphasizing the need for long read sequencing in crop species to identify novel variation. Furthermore, we highlight SVs and SNPs occurring in genes and pathways with known associations to critical bioenergy-related phenotypes and characterize the landscape of genetic differences between sweet and cellulosic genotypes.

## Introduction

Sorghum (*Sorghum bicolor* (L.) Moench) is a versatile, adaptable, and widely grown cereal crop that is valued for its efficiency, drought tolerance, and ability to grow in marginalized soils (Wayne Smith and Frederiksen, 2000). Present-day genotypes exhibit extensive genetic, phenotypic, morphological, and physiological diversity which stems both from their historical spread and modern breeding efforts aimed at optimizing sorghum for different end uses. With its wealth of naturally occurring genetic diversity and advantageous traits, sorghum has enormous value as a sustainable, fast-growing, and high-yielding bioenergy crop (Calviño and Messing, 2012).

Currently, sorghum is classified into four major ideotypes: grain, sweet, cellulosic, and forage. All of these types can be used in different bioenergy production methods (Wu *et al*., 2010), but to fully capitalize on their potential, it is essential to gain a better understanding of the genomic changes driving traits related to yield, carbon partitioning, and local adaptation. However, these types of traits are often difficult to dissect due to the nature of their underlying genetic architecture (Brachi, Morris and Borevitz, 2011), which can involve hundreds to thousands of genes and complex mutations that are not easily captured by short-read sequencing.

Structural genomic mutations are an important source of variation in many species, and can play key roles in phenotypic diversification and evolution. Advances in sequencing technology, especially the advent of high-throughput long-read sequencing, have made the detection of structural variants feasible in many plant species where these types of changes were previously uncharacterized. More recently, there has also been a surge in the generation of pan-genomic data for a number of important crop species, which has offered exciting new insights into the extensive diversity of these plants and the potential influence of complex structural mutations on agronomically important phenotypes (Golicz, Batley and Edwards, 2016; Zhang *et al*., 2019; Danilevicz *et al*., 2020; Zhou *et al*., 2020, 2022; Della Coletta *et al*., 2021; Hufford *et al*., 2021; Li *et al*., 2021).

Previous genomic work in sorghum has linked structural mutations to a number of key traits including dwarfing (Multani *et al*., 2003), juicy stalks (Zhang *et al*., 2018), chilling tolerance (Wu *et al*., 2019), and flowering time (Li *et al*., 2018). A whole-genome comparison of the sweet sorghum genotype ‘Rio’ with ‘BTx623,’ (a short-statured, early maturing grain sorghum) found hundreds of gene presence/absence variations (PAVs), several of which occurred among known sucrose transporters (Cooper *et al*., 2019). Furthermore, a genome-wide association study (GWAS) exploring the genetic architecture of bioenergy-related traits found that a large deletion in a sorghum-specific iron transporter was linked to stalk sugar accumulation (Brenton *et al*., 2016, 2020). Most recently, we undertook a broad survey of genome-wide deletions in a panel of nearly 350 diverse sorghum accessions, and found large deletions in multiple genes related to biotic and abiotic stress responses that were unique to particular geographic origins, and appeared to play a role in local adaptation (Songsomboon *et al*., 2021).

Taken together, these results suggest that unraveling complex traits in sorghum and other crops will require a comprehensive picture of both structural and single nucleotide mutations. In this study, we have expanded on the recently published Carbon-Partitioning Nested Association Mapping (CP-NAM) population that was developed and publicly released as a key genetic resource for the characterization and improvement of sorghum for multiple different end uses (Boatwright *et al*.,2021, 2022; Kumar *et al*., 2022). We generated high-quality *de novo* genome assemblies for 10 of the CP-NAM parents and used these genomes to identify millions of novel variants, including a number of large structural variants (SVs) occurring in genes or pathways that could be essential for optimizing sorghum as a bioenergy feedstock.

## Materials and Methods

### Sample Collection and Sequencing

Seeds for each genotype were ordered from the U.S. Department of Agriculture’s Germplasm Resource Information Network (GRIN)(https://www.ars-grin.gov/) and grown in the greenhouses at the North Carolina Research Campus (NCRC) in Kannapolis, NC. High-molecular-weight DNA was extracted from each sample using a modified high-salt CTAB extraction protocol (Inglis *et al*., 2018). Purified DNA was sent to the David H. Murdock Research Institute (DHRMI) for quality control, library preparation, and sequencing on a PacBio Sequel I system.

### De novo Assembly

Raw subreads for each genotype were combined and converted to FASTQ format using the `bam2fastx` toolkit from PacBio. Reads were then corrected, trimmed, and assembled using Canu(v2.1.1) (Koren *et al*., 2017). For one of the genotypes, ‘Grassl’, Canu failed to produce contigs due to reduced read coverage after trimming, so the final assembly was instead produced using Flye(v2.9) with the Canu corrected reads (Kolmogorov *et al*., 2019).

The resulting contigs for all genotypes were scaffolded into chromosomes using RagTag (v2.1.0)(Alonge *et al*., 2021) and the parameters ‘-r -g 1 -m 10000000’. Contigs were ordered based on their alignment to the BTx623 v3.1 reference genome (Paterson *et al*., 2009) with minimap2 (Li, 2018). RagTag was run *without* the correction step to avoid unnecessary fragmentation of the contigs and unplaced contigs were discarded. Assembled genome metrics were assessed both before and after scaffolding using QUAST(5.2.0) (Gurevich *et al*., 2013).

### Annotation

Protein and non-coding genes were annotated by building a pan-gene working set using representative pan-gene models selected from a comparative analysis of gene family trees from 18 Sorghum genomes (McCormick *et al*. 2018; Deschamps *et al*. 2018; Cooper *et al*. 2019; Wang *et al*. 2021; Tao *et al*. 2021) sourced from SorghumBase(https://www.sorghumbase.org/). This pan-gene representative was propagated onto the 10 sorghum genome assemblies using Liftoff (v1.6.3)(Shumate and Salzberg 2021) with default parameters. The gene structures were updated with available transcriptome evidence from Btx623 using PASA (v2.4.1)(Haas *et al*. 2003). Additional improvements to structural annotations were done in PASA using full length sequenced cDNAs and sorghum ESTs downloaded from NCBI using the query (EST[Keyword]) AND sorghum[Organism]. The working set was assigned Annotation Edit Distance(AED) scores using MAKER-P (v3.0)(Campbell *et al*. 2014) and transcripts with AED score < 1 were classified as protein coding. Those with AED=1 were further filtered to keep any non-BTx623 based models with a minimum protein length of 50 amino acids and a complete CDS as protein coding. The remaining models with AED=1 were classified as non-coding. Gene ID assignment was made as per the existing nomenclature schema established for Sorghum reference genomes(McCormick *et al*. 2018).

On average, approximately 55 thousand working sets of models were generated for each sorghum line, out of which an average of 41 thousand were coding and roughly 13 thousand were non-coding (Supplementary Table 1). More than half (61%) of the protein coding models mapped to a BTx623 reference gene, along with 23% of the non-coding models (Supplementary Figure 1). Functional domain identification was completed with InterProScan (v5.38-76.0) (Jones *et al*. 2014). TRaCE (Olson and Ware 2020) was used to assign canonical transcripts based on domain coverage, protein length, and similarity to transcripts assembled by Stringtie. Finally, the protein coding annotations were imported to Ensembl core databases, verified, and validated for translation using the Ensembl API (Stabenau *et al*. 2004).

In order to assign gene ages, protein sequences were aligned to the canonical translations of gene models from *Zea mays, Oryza sativa, Brachypodium distachyon*, and *Arabidopsis thaliana* obtained from Gramene release 62 (Tello-Ruiz *et al*. 2020) using USEARCH v11.0.667_i86linux32 (Edgar 2010).If there was a hit with minimum sequence identity of 50% (-id 0.5) to an *Arabidopsis* protein, the gene was classified as being from Viridiplanteae, if there was a hit to rice the gene was classified as Poaceae, and if a hit was to maize the gene was classified as Andropogoneae. If there were no hits then the gene was classified as sorghum specific.

### Repeat Analysis

Transposable elements (TEs) were identified and annotated in each genome using EDTA (Ou *et al*., 2019). TE-greedy-nester (Lexa *et al*., 2020) was used to further annotate both complete and fragmented Long Terminal Repeat (LTR) retrotransposons. Sequence divergence in the LTR regions was used to estimate retrotransposon age (SanMiguel *et al*., 1998; Jedlicka, Lexa and Kejnovsky, 2020). The left and right LTR sequences were extracted from the assembled genomes using the coordinates reported by TE-greedy-nester and the `getfasta` tool from the BEDTools package(v2.29.0) (Quinlan and Hall, 2010). For each TE, the two LTR sequences were aligned using Clustal-W (Thompson, Higgins and Gibson, 1994) as implemented in the R package msa (Bodenhofer *et al*., 2015). Genetic distance was calculated based on the K80 model using the `dist.dna` function in the R package phangorn (Schliep, 2011). The time of divergence was calculated based on the equation T=K/(2 * r) (Bowen and McDonald, 2001), where T is the time of divergence, K is the genetic distance, and r is the substitution rate. A value of 0.013 mutations per million years was used for r, consistent with the molecular clock rate for LTRs estimated in rice (Ma and Bennetzen, 2004).

### Variant Calling

Filtered and scaffolded reads were realigned to the BTx623 reference genome using the `nucmer` program from the MUMmer(v4.0) package (Delcher, Salzberg and Phillippy, 2003; Marçais *et al*.,2018) with the following parameters ‘-c 100 -b 500 -l 50’. Alignments were filtered using the `delta-filter` program from the MUMmer package with the parameters ‘-m -i 90 -l 100’ and converted to coordinate files using `show-coords` with the parameters ‘-THrd’. Variants were then called using Syri(v1.6)(Goel *et al*., 2019).

Individual Syri VCF files were split by variant type (SNPs, Deletions, Insertions, Inversions, and Translocations) resulting in separate files for each variant type for each genotype. Insertions or deletions smaller than 50 bp were classified as small indels while those equal to or larger than 50 bp were classified as SVs. More complex SV types that could not be validated with raw reads were not considered for further analysis.

The Syri program produces a nonstandard VCF format which includes information on variants from overlapping syntenic blocks. This can result in duplicated variants and fragmented insertions that must be addressed before subsequent analysis with downstream tools. Duplicates of existing variants were removed for all variant types, and fragmented insertions were combined into single variants (Supplementary Figure 2). These processed variant files were then zipped and indexed using `bgzip` and `tabix` (Li *et al*., 2009) and then merged across genotypes using the merge function from the bcftools package with the parameters ‘-0 -I ‘ChrB:join,Parent:join,DupType:join,modified:join’ -O v’. This resulted in one variant file for each type of variant that included the genotypes for all individuals. Insertions, deletions, and SNPs were then annotated using SIFT (v2.4)(Vaser *et al*.,2016) and the BTx623 version 3.1.1 annotation to identify overlap with genes for insertions and deletions and missense prediction for single nucleotide variants.

### Phylogeny

Gene PAVs were called from pan-gene lift-off annotation information using custom python scripts. PAVs for each genotype were encoded as a binary vector (with 0 indicating gene absence, and 1 indicating presence). Distance between genotypes was then calculated using the dist() function from the stats(v3.6.2) package in R using the Jaccard distance, and a phylogenetic tree was constructed using the NJ() function from the phangorn package.

### Gene Ontology Analysis

Gene ontology (GO) terms for genes affected by large insertions and deletions or nonsynonymous SNPs were curated from the publicly available annotation information file associated with BTx623 v3.1.1 in phytozome (https://phytozome-next.jgi.doe.gov/). GO enrichment analysis was performed using the R package topGO(v1.0) (Alexa and Rahnenfuhrer, 2016).The classic Fisher’s Test was used to assess significance of enriched terms, and terms with a p-value <0.05 were considered significant and kept for further analysis. Redundant and highly similar GO terms were defined and reduced based on semantic similarity using the R packages AnnotationForge (Carlson and Pages,2022) and rrvgo (Sayols, 2020).

## Results

### Assembly Quality and Characteristics

To capture the genetic diversity of bioenergy sorghum, we sequenced the parents of the previously established CP-NAM population, which included globally diverse genotypes representative of sweet, cellulosic, grain and forage type bioenergy sorghums (Boatwright *et al*., 2021)(Table 1). The initial contig-level assemblies showed a range of N50 values, with the lowest being 176 kb and the highest at over 3 Mbp (Supplementary Table 2). The three sweet genotypes in particular had a higher number of raw reads and more contiguous assemblies than the other types (Figures 1A and 1B), most likely as a result of differences in the effectiveness of the extraction protocol. After scaffolding and filtering unplaced contigs, all 10 genotypes showed similar levels of high contiguity, with final assembly sizes that were 90-98% the size of the BTx623 reference genome and over 90% of known BTx623 genes contained within the scaffolds (Figures 1C and 1D).

**Figure 1.**
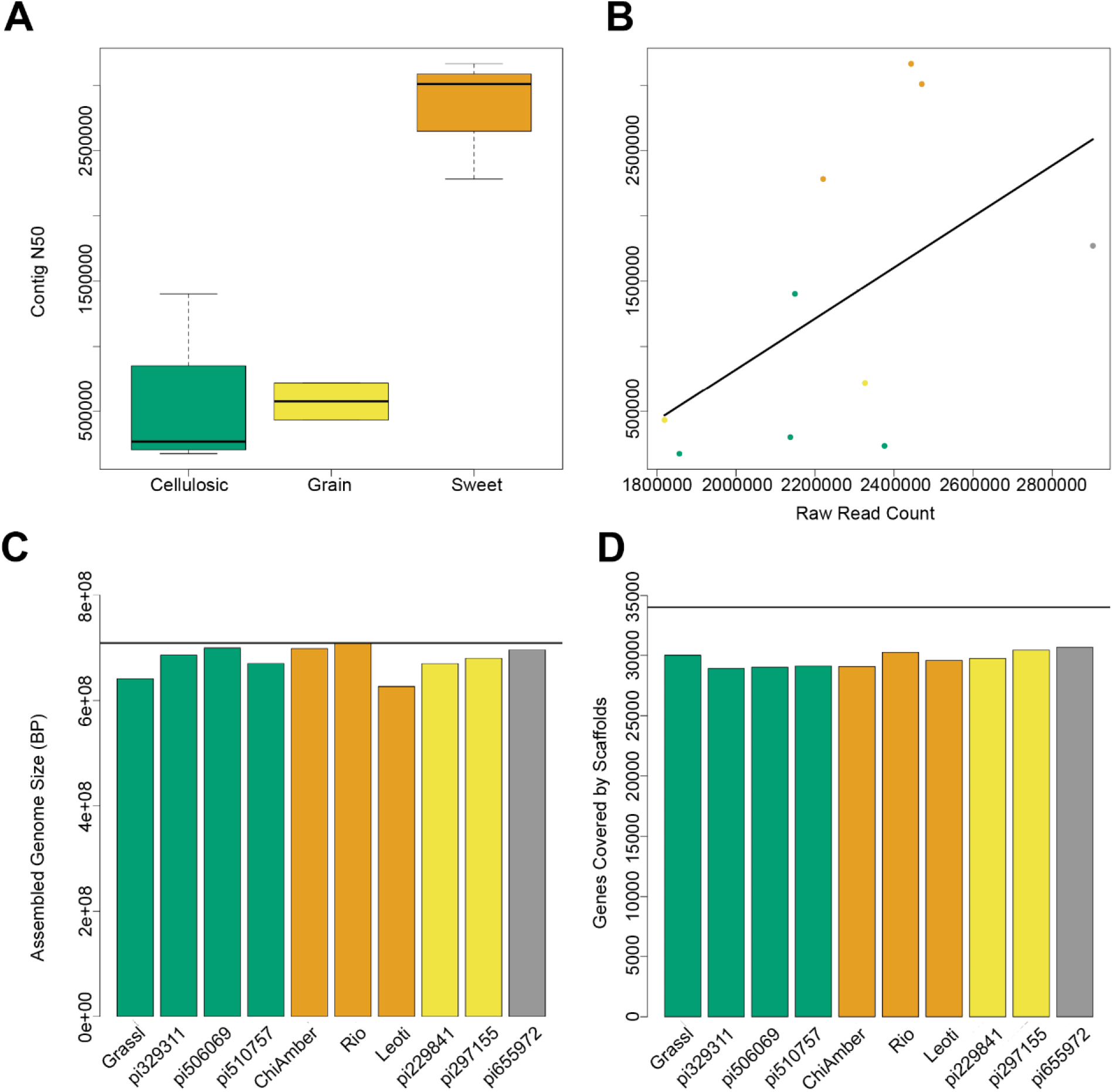
Assembly metrics for 10 sorghum genotypes. A) Contig N50 levels for different ideotypes show higher contiguity for sweet genotypes. B) Raw read counts prior to assembly are highly correlated with contig N50, and sweet genotypes (orange) have higher read counts than cellulosic (green) or grain (yellow) genotypes. C) Assembled genome size after scaffolding and filtering for each genotype shows that despite differences in mean contig size, the final assemblies for both sweet and non-sweet types are very close to the expected reference genome size (horizontal black line). D) The number of BTx623 genes contained within the final scaffolds is very similar across all genotypes regardless of type.

**Table 1.** Genotype Origins, Races, and Types.

| Name | Alternate ID | Race | Origin | Type |
| --- | --- | --- | --- | --- |
| Grassl | PI 154844 | Caudatum | Uganda | Sweet & Cellulosic |
| PI 329311 | IS 11069 | Durra | Ethiopia | Cellulosic |
| PI 506069 | Mbonou | Guinea-bicolor | Togo | Cellulosic |
| PI 510757 | AP79-714 | Durra | Cameroon | Cellulosic |
| Chinese Amber | PI 22913 | Bicolor | China | Sweet |
| Rio | PI 563295 | Durra-caudatum | USA | Sweet |
| Leoti | PI 586454 | Kafir-bicolor | Hungary | Sweet |
| PI 229841 | IS 2382 | Kafir | South Africa | Grain |
| PI 297155 | IS 13633 | Kafir | Uganda | Grain, Forage |
| PI 655972 | Pink Kafir | Kafir | USA | Forage |

Information adapted from GRIN and (Boatwright *et al.*, 2021).

### Gene Annotation

Genes shared across deeper evolutionary time scales were more conserved than sorghum-specific genes (Figure 2). The sweet genotypes show slightly more conserved genes when compared to other genotypes (Figure 2). Around 36.69 percent of genes were found to be core to all genotypes, 50.32 percent were shell genes (present in more than one genome, but not all of the genomes), and 12.99 percent were found to be cloud genes (unique to a single genome) (Supplementary Figure 3). Of shell genes identified, 44 and 45 were identified to be exclusive to all sweet and all non-sweet genotypes respectively.

**Figure 2.**
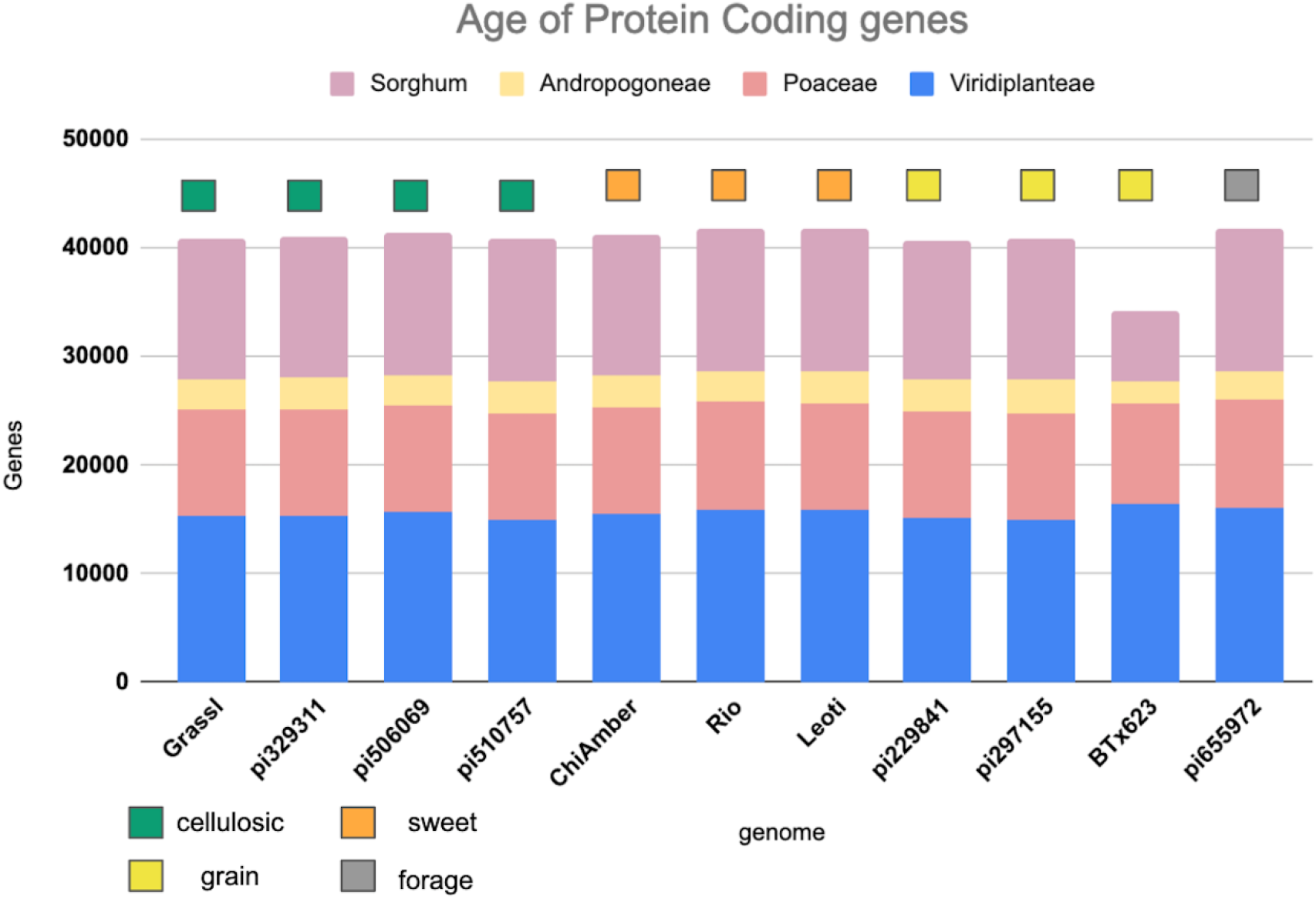
Age of protein coding genes among the sorghum lines based on minimum sequence identity. Bar color indicates the level of phylogenetic conservation, with blue indicating genes conserved across monocots and dicots; peach indicating the proportion of genes shared among the grasses; yellow indicating the proportion of genes shared between sorghum and maize, and light purple representing the proportion of sorghum-specific genes.

### Genomic Landscape of Variation

Over 10.5 million single nucleotide variants were called across the 10 genomes, as well as over 7.4 million small indels and over 24 thousand large structural variants (insertions and deletions ≥ 50 bp) (Figure 3, Tables 2 and 3). Well over half (~65%) of these variants were defined as cloud variation (Table 3), while the remaining variants were mostly shell. Only a small handful of core variants were present in all of the genotypes except the BTx623 reference. Phylogenetic relationships were inferred using gene presence/absence to estimate genetic distance (Supplementary Figure 4), demonstrating that sweet, cellulosic, and grain genotypes come from separate clades within the category of bioenergy-type sorghum.

**Figure 3.**
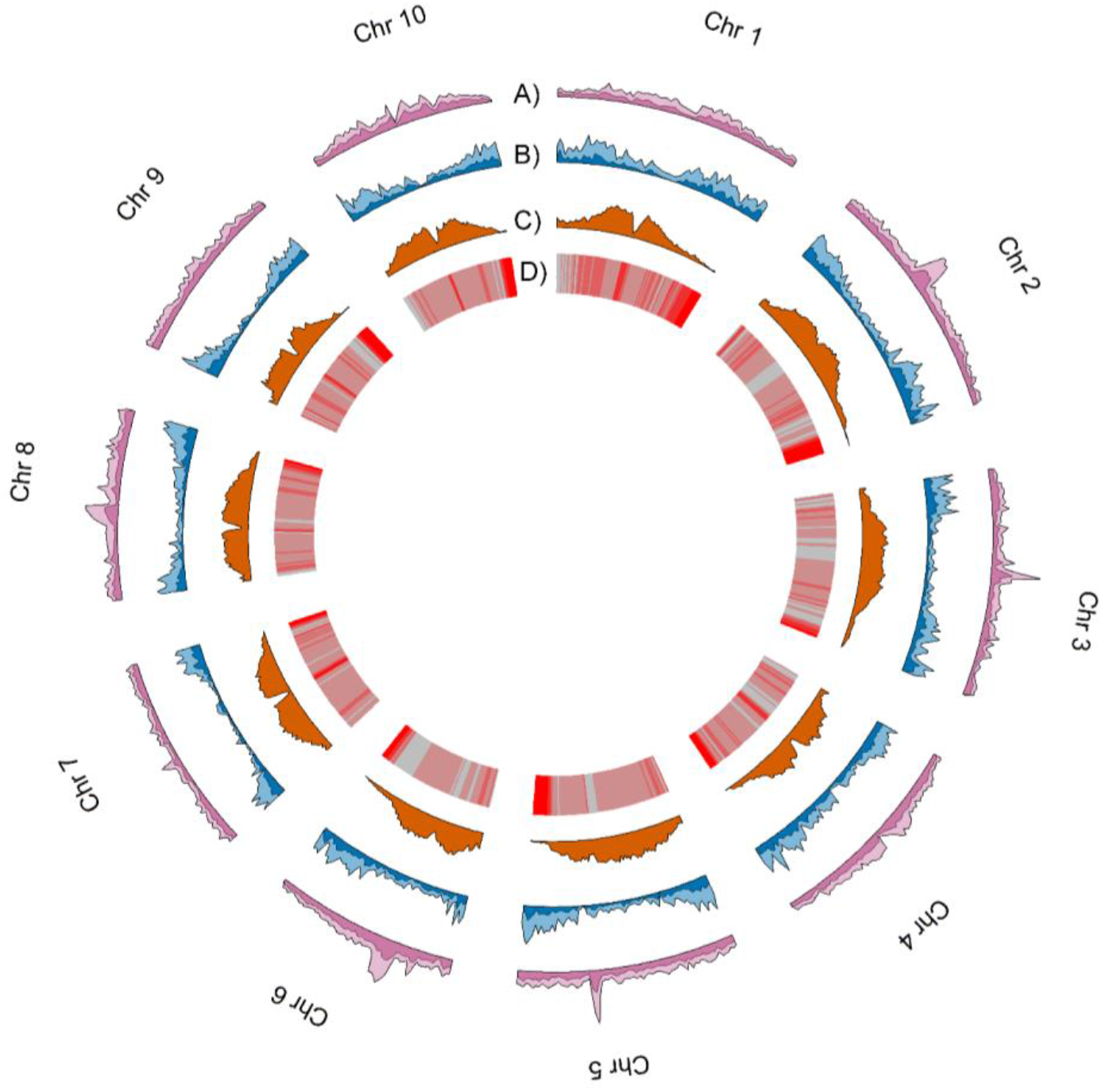
Genomic landscape of variation averaged across the 10 genomes. Density estimates in tracks A-C were performed in 1Mb non-overlapping sliding windows. A) and B) respectively show average SNP density and average SV density, with lighter colors indicating cloud variants and darker colors indicating shell and core variants. C) shows the average TE density, and D) shows TE age averaged across 1Mb sliding windows. Red indicates younger TEs while gray indicates older.

**Table 2.**
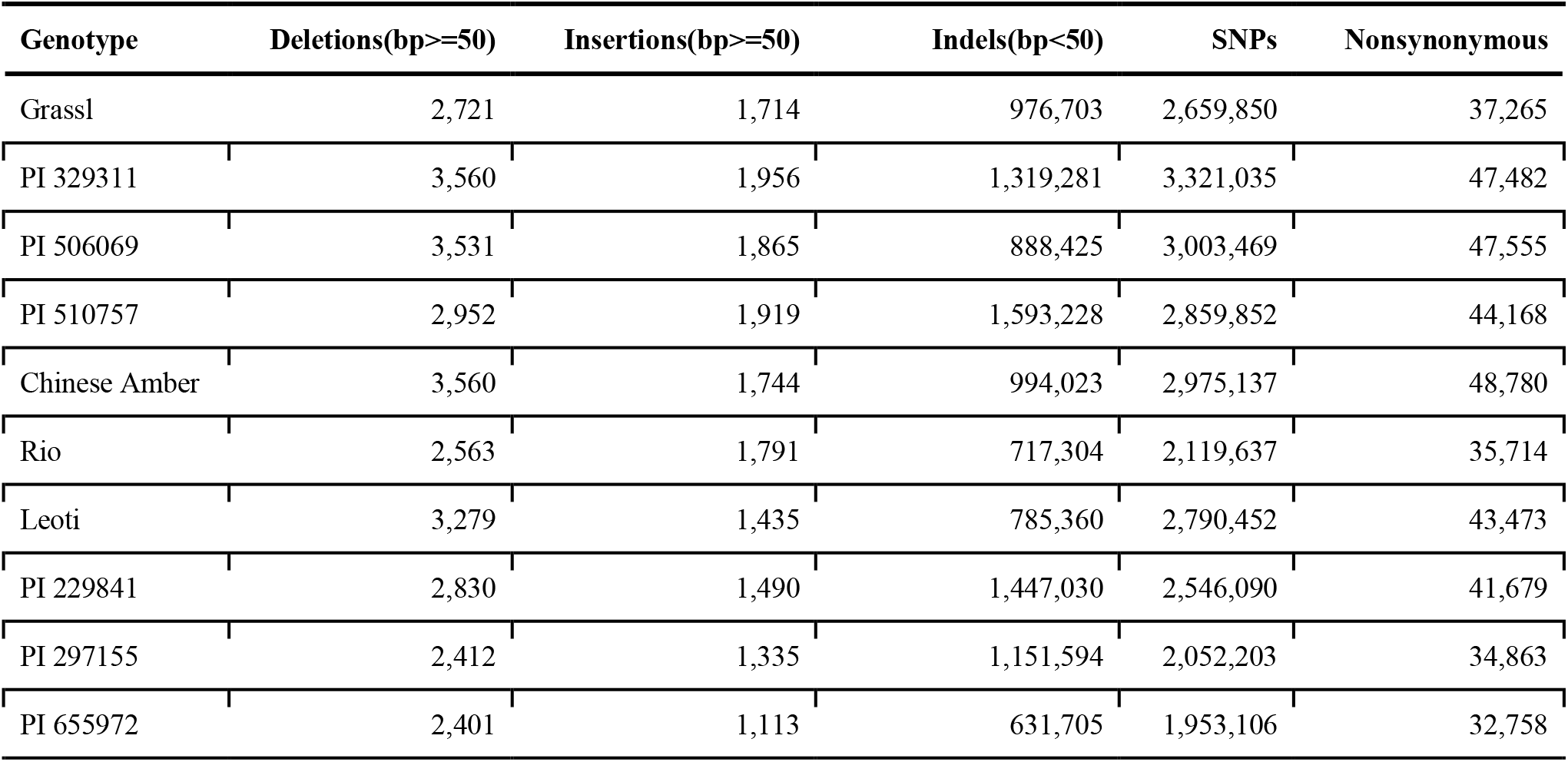
Variants found in each NAM parent genotype.

**Table 3.**
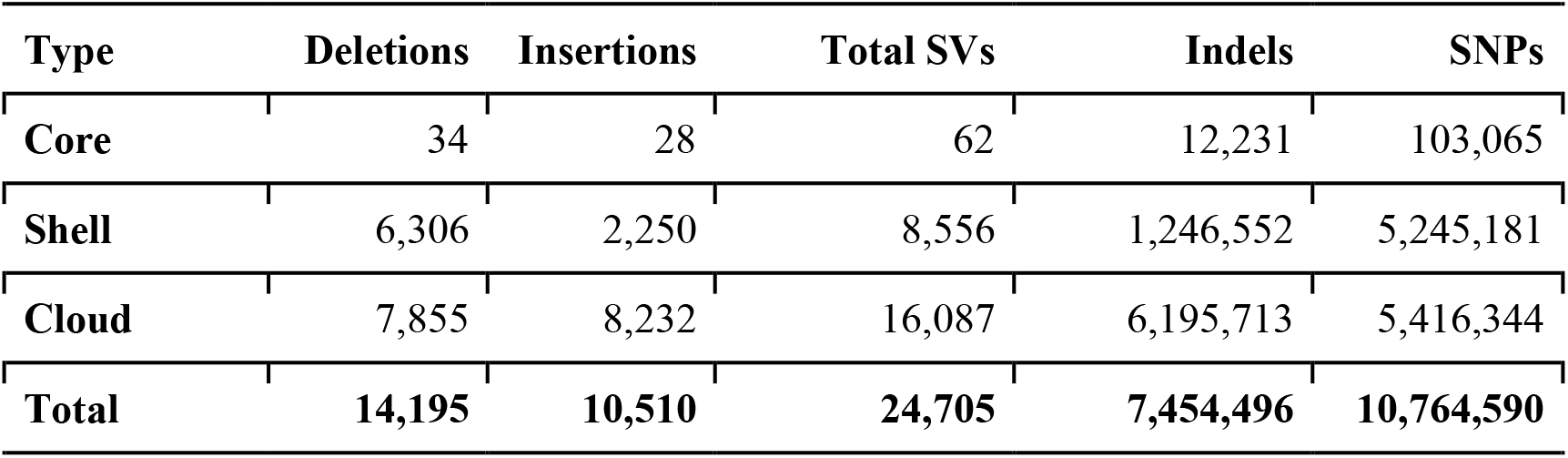
Core vs. Shell vs. Cloud variants

### Genes Affected by Structural Variants and SNPs

There were a total of 171,000 SNPs that were found to be both located in genic regions and encoding nonsynonymous variants, and more than 2.5 thousand large SVs present in genic regions. GO enrichment analyses of affected genes revealed that SNPs and SVs tended to impact distinct categories of genes (Figure 4), with protein phosphorylation being the only significant category to appear in both datasets.

**Figure 4.**
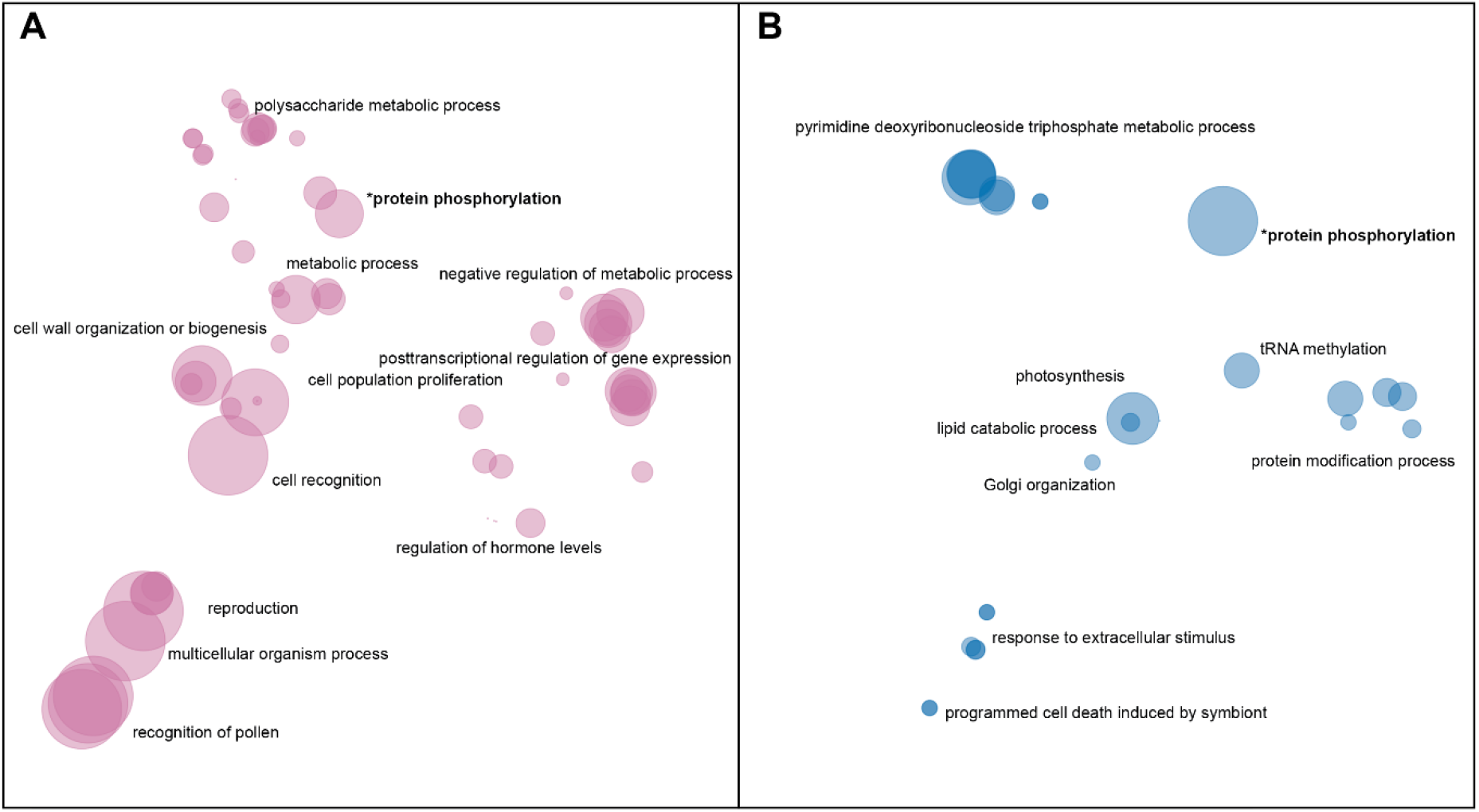
Enriched GO terms for genes impacted by A) nonsynonymous SNPs and B) large SVs. GO terms in each dataset were clustered and plotted based on semantic similarity as described in the Materials and Methods. Circle size is proportional to p-value, with larger circles indicating more significant terms.

In addition to protein phosphorylation, genes impacted by large insertions or deletions showed enrichment in GO categories related to Golgi vesicle transport, photosynthesis, nucleoside metabolism, protein modifications, and programmed cell death (Figure 4B). Nonsynonymous SNPs, on the other hand, were enriched in genes involved in pollen-pistil interactions, cell wall biogenesis, cell proliferation, posttranscriptional regulation and polysaccharide metabolism (Figure 4A).

### Repeat Analysis

Overall the TE composition was highly similar across all 10 genotypes (Figure 5 and 3), with the LTR-Gypsy superfamily comprising the majority of elements. The age analysis revealed an abundance of younger TEs, with a mean age of 1.28 million years old along with a high frequency of very young TEs approximately 0.1 million years old and very few old TEs (6-8 million years) (Figure 5; Supplementary Figure 5). Most (97.5%) of the TEs were non-nested, with TE-greedy-nester reporting the presence of only a handful (2.5%) of nested TEs. The overall distribution of TE age followed a similar pattern across all of the genotypes, with younger TEs being randomly distributed throughout the genome (Figure 3, Supplementary Figure 6A-J) as previously observed by (Paterson *et al*., 2009).

**Figure 5.**
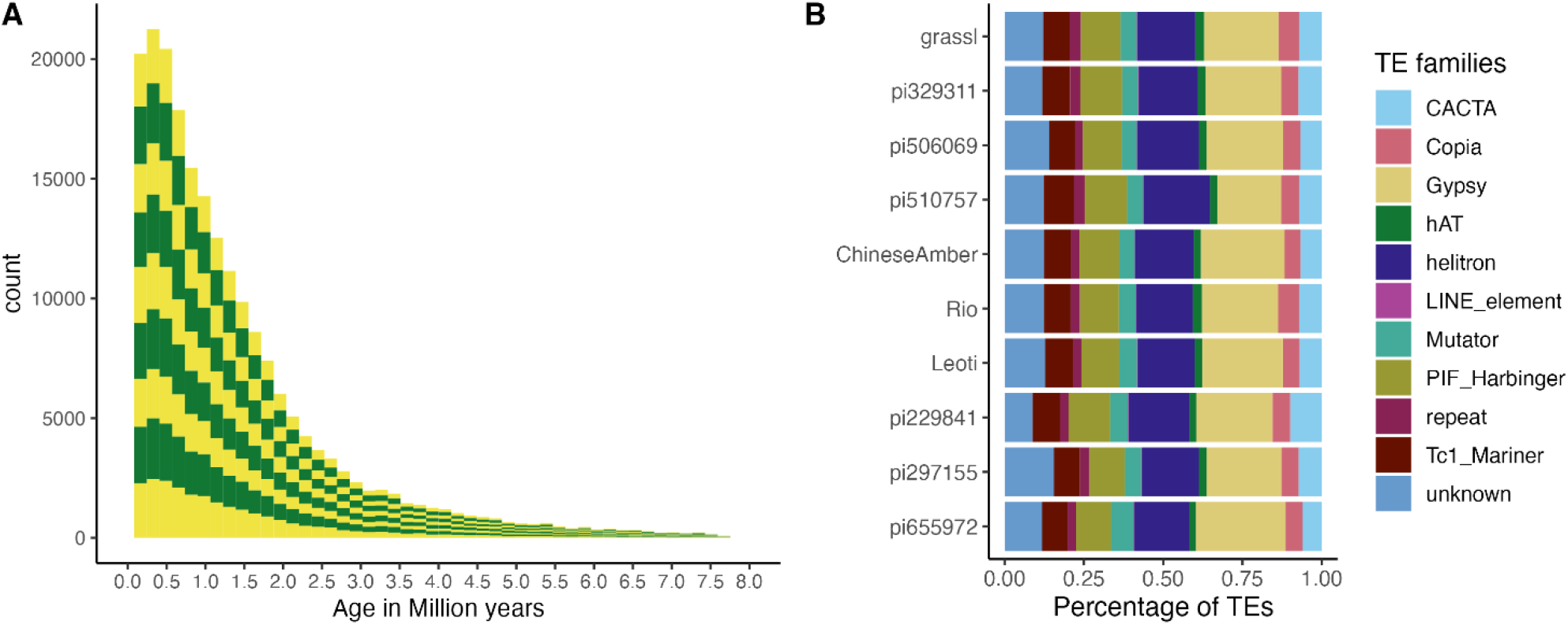
TE age and composition. A) The distribution of TE counts by age across all genotypes. Alternating colors indicate different genotypes, with the Y-axis labeling the number of TEs and the X-axis labeling their age in millions of years. B) The proportion of superfamilies of TEs based on average counts of each superfamily across all genomes.

### Differences in Sweet and Non-Sweet Genotypes

Structural variants that were present in all three sweet genotypes (Leoti, ChineseAmber, and Rio) but either absent from or rare among non-sweet genotypes, were significantly enriched among genes with functions related to metal ion transport, in particular iron ion transport, as well as genes involved in oxidative stress response, cell cycle arrest, and phosphatidylserine biosynthetic processes. Conversely, variants found only in all of the non-sweet genotypes tended to impact very different categories of genes, such as those involved in glycolytic processes, cytochrome assembly, and both RNA and DNA regulation (Figure 6).

**Figure 6.**
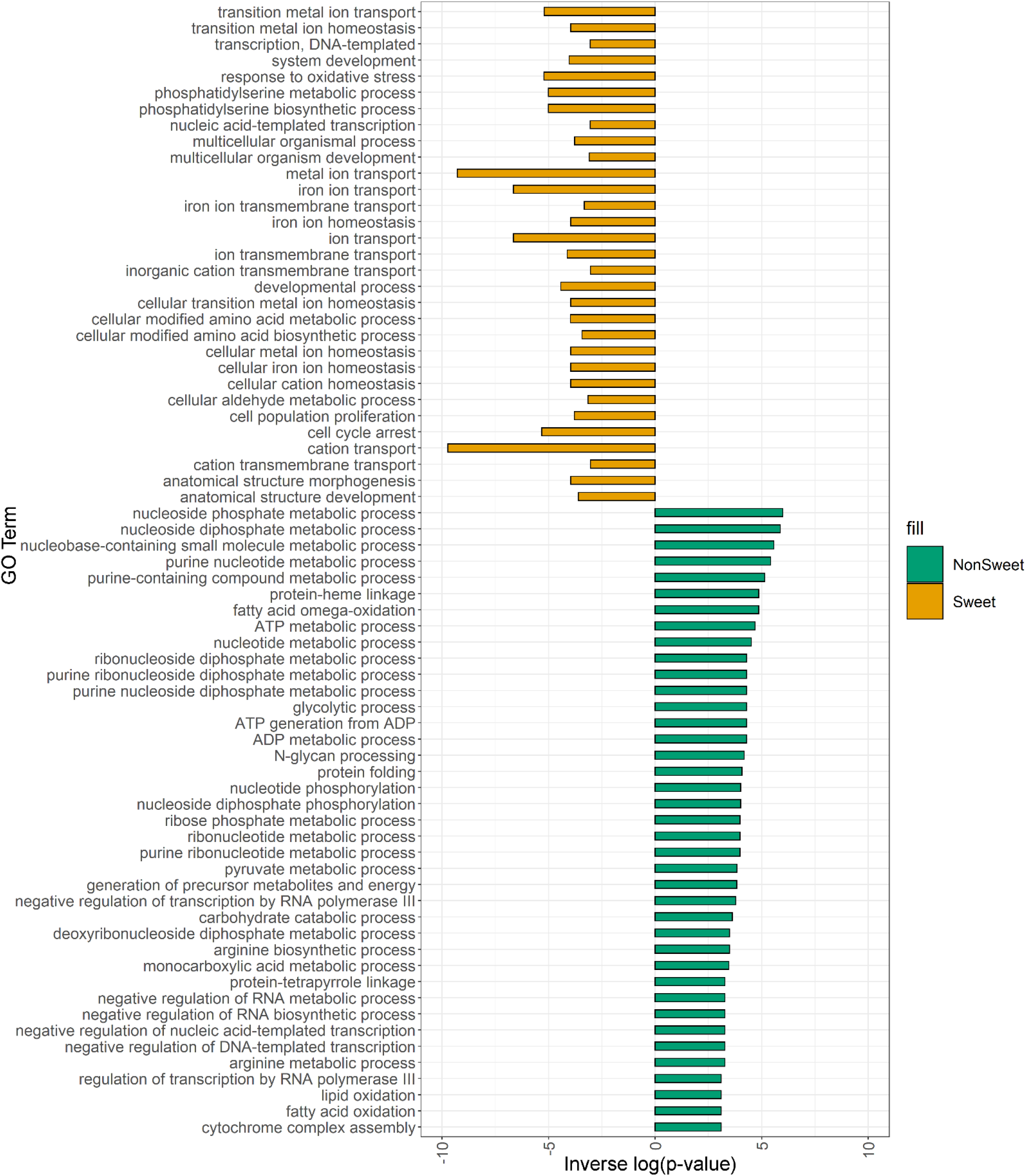
Enriched GO terms for genes impacted by SVs and Indels in both Non-Sweet and Sweet Genotypes. Orange bars indicated gene categories in Sweet genotypes that were significantly impacted (p<0.05). Green bars indicated gene categories in Non-Sweet genotypes that were significantly impacted (p<0.05). The length of each bar corresponds to significance (−log(p-value)).

## Discussion

Unraveling the molecular mechanisms controlling complex traits such as carbon partitioning, yield, and stress response is an essential step for crop improvement efforts aimed at creating effective and sustainable bioenergy feedstocks for the future. However, not only do these types of traits often involve changes in large numbers of genes, but an ever-increasing number of pan-genomics studies in crop plants have demonstrated that these changes can encompass complex structural mutations in addition to SNPs (Cooper *et al*., 2019; Zhang *et al*., 2019; Brenton *et al*., 2020; Zhou *et al*., 2020, 2022; Hufford *et al*., 2021; Songsomboon *et al*., 2021). Therefore, the development of multiple reference-quality genomes within crop species is critical to the exploration of complex genetic architectures and has clear benefits when compared to a single reference genome, especially in the case of larger structural variants(Della Coletta *et al*., 2021). By *de novo* assembling 10 new high-quality genomes for the parents of the CP-NAM population (Boatwright *et al*., 2022), we have been able to uncover millions of novel variants, including thousands of large insertions and deletions.

Importantly, we found that SVs within coding regions impacted different types of genes compared to SNPs, highlighting the importance of incorporating both into future trait mapping studies. Many nonsynonymous SNPs that were segregating among the genotypes occurred in gene categories that have previously been linked to carbon allocation in sorghum and other closely related species. For instance, protein phosphorylation induces key signaling cascades in plants that control a variety of processes, and protein kinases have been shown to be highly differentially expressed in both sweet sorghum (Cooper *et al*., 2019) and sugarcane (Waclawovsky *et al*., 2010) during stem sugar accumulation. Similarly, genes involved in the regulation of plant hormones such as auxin were also enriched for non-coding SNPs, and these pathways are known to be essential for vegetative plant growth and stem elongation, both of which are key phenotypes for biomass accumulation (Kebrom, McKinley and Mullet, 2017).

Like SNPs, gene-impacting SVs were also found to affect many genes related to protein phosphorylation; in fact, this was the top category among genes containing large variants. But other categories enriched for high-impact insertions and deletions were distinct from the SNP dataset, and contained many genes involved in pathways related to both abiotic and biotic stress responses, which has been observed before in diverse bioenergy sorghums (Songsomboon *et al*., 2021). Additionally our study identified structural variants affecting genes involved in tRNA nucleoside modifications, programmed cell death in response to symbionts, and photosynthetic light response, all of which were previously identified by other studies as GO terms of interest in relation to sorghum stress response (Ortiz, Hu and Salas Fernandez, 2017; Wang *et al*., 2017).

SVs strictly occurring in either sweet or non-sweet genotypes also offer unique insights into the differences between these types that could be key to dissecting differences in carbon allocation in sorghum. Of particular interest is the fact that SVs restricted to sweet sorghum genotypes affected many genes related to metal metabolism and iron transport. This connection between iron transport and sugar accumulation has been observed in other comparative genomic studies of sorghum (Brenton *et al*., 2016, 2020; Cooper *et al*., 2019), and appears to be a key factor distinguishing sweet sorghums from both cellulosic and grain types.

Over a third of protein coding genes and over 75 percent of noncoding genes annotated in this study did not map back to the Btx623 reference genome. With a growing number of studies illustrating the importance of noncoding DNA and RNA as potential regulatory elements (Waititu et al. 2020), it is evident that large pan-genome annotations are vital in quickly identifying and annotating potential regulatory ‘pseudo-genes’ as well as protein coding genes that are divergent from the common reference. Previous pan-genome studies in sorghum and maize have identified high levels of gene content variation, with 53-64 percent of genes identified as non-core (Tao *et al*., 2021;Ruperao *et al*.,2021;Hufford *et al*., 2021). We corroborate these findings with about 63 percent of our genes being identified as either shell or cloud to our population, despite this particular population lacking wild representation, indicating relatively high amounts of latent variation, even among domesticated varieties of sorghum.

Taken together, our results demonstrate the value of exploring genome-wide patterns of both SNPs and larger structural variants to gain new insights into the genetic architectures of complex and agronomically important traits. To advance both sorghum breeding efforts and our understanding of crop plant evolution, we have generated this new extensive dataset that is publicly available through SorghumBase (Gladman *et al*., 2022) and which can be readily integrated into an already valuable genetic resource for future mapping studies.

## Nomenclature

CP-NAM: Carbon Partitioning Nested Association Mapping
SV: Structural Variant
SNP: Single Nucleotide Polymorphism
TE: Transposable Element
LTR: Long Terminal Repeat
GO: Gene Ontology

## Conflict of Interest

The authors declare that the research was conducted in the absence of any commercial or financial relationships that could be construed as a potential conflict of interest.

## Author Contributions

WGV: Writing, variant analysis, created figures and tables, performed scaffolding.

KK: Performed TE analysis, Alignments, and Variant calling. Wrote corresponding methods sections.

LCA: Aided in scripting of figure creation and filtering of variants.

KS: Growing and DNA Extraction of plant material.

CP: Aided in genome assembly.

KC, ZL, AO: Gene and transposable element annotations.

DW: Experimental design, writing.

ZWB: designed CP-NAM population, provided genetic materials

JLB: development and release of CP-NAMs

EAC: Writing, created figures, conceived the project, advised, and helped direct analysis.

## Funding

This research was supported by startup funds from UNC Charlotte and the United States Department of Agriculture grant USDA-ARS 8062-21000-041-00D.

## Acknowledgments

The authors would like to thank S. Kresovich and M. Myers for providing plant materials, J. Lotito and N.C. State for providing and overseeing the greenhouse and growth chambers facilities at the N.C. Research Campus, and the DHMRI Genomics Core for providing sequencing services. The authors would also like to acknowledge the University Research Computing team at UNC Charlotte and S. Blanchard for providing essential IT support and resources.

## Data Availability Statement

Assembled Genomes are publicly available on https://www.sorghumbase.org/. Gene data hosted at https://ftp.sorqhumbase.org/Voelker_et_al_2022/.Raw data and genome assemblies are available at the European Nucleotide Archive under the project ID: PRJEB55613

## Notes

### Competing Interest Statement

The authors have declared no competing interest.

